# Conflict of evolutionary interests between plants and pollinators revealed through functional exploration of flower morphospace

**DOI:** 10.1101/370700

**Authors:** Foen Peng, Eric O. Campos, Joseph Garret Sullivan, Nathan Berry, Bo Bin Song, Thomas L. Daniel, H. D. Bradshaw

## Abstract

The explosive evolutionary diversification of flowering plants traditionally is attributed to the coevolution of plants and their animal pollinators. Plant-pollinator interactions are held as classical examples of mutualisms – beneficial to both parties – in spite of the fact that most other cases of rapid coevolution are the result of conflicts of interest (*e.g.*, predator-prey, host-parasite, sexual conflict, competition for resources). Could the co-diversification of plants and their pollinators be driven by conflict rather than by mutualism? To address this question, we employed the theoretical morphospace paradigm using a combination of 3D printing, electronic sensing, and machine vision technologies to determine the influence of two flower morphological features (corolla curvature and nectary diameter) on the fitness of both parties: the artificial flower and its hawkmoth pollinator. We found that the two parties have almost opposite interests in corolla curvature evolution, with non-overlapping fitness peaks in flower morphospace, suggesting that the evolutionary radiation of flowering plants and their pollinators could be the result of conflict instead of mutualism.

## Introduction

Flowering plants (angiosperms) are the most diverse lineage in the plant kingdom. Their initial major diversification happened in the early Cretaceous (about 130 - 190 Mya), which was accompanied by the co-radiation of pollinating insects [1,2]. This rapid co-diversification process has been attributed to mutual adaptations for biotic pollination. At least a quarter of divergence events in flowering plants are associated with pollinator shifts [3].

Traditionally, plant-pollinator interactions are considered to be classical examples of mutualism – increasing the fitness of both parties [4]. Pollinators provide plants with pollen transport services leading directly to offspring production, while plants provide pollinators with many types of rewards, such as energy-rich nectar [5], shelter [6], or thermoregulation [7], that enhance the survival, growth, and reproduction of the pollinators.

However, most examples of rapid coevolutionary diversification are the result of conflicts of interest rather than mutualisms [8]. For example, food competition drives stickleback populations to diverge into a “limnetic” species and a “benthic” species [9]. Similarly, the arms race between plant (prey) chemical defenses and caterpillar (predator) counter-defenses promotes diversification in both groups [10]. Host-parasite [11] and male-female conflicts [12] likewise increase the rates of coevolutionary diversification.

In contrast, mutualistic coevolution assumes that each party maximizes its own fitness only when its phenotype matches the needs of its partner. These matching phenotypes should be maintained by stabilizing selection, thus hindering diversification [8]. The orthodox view of the plant-pollinator relationship as a mutualism cannot easily explain the rapid diversification of angiosperms and insects. Therefore, we test the counterintuitive hypothesis that there is a conflict of interest between plants and their pollinators.

Flower morphology is key to understanding plant-pollinator interactions, which is exemplified by Darwin’s famous prediction of the existence of a hawkmoth species with a long proboscis to match the extraordinarily long nectar spur of a Malagasy orchid [13]. Previous studies have shown that many flower morphological traits, such as anther position [14,15], corolla tube length [16], corolla width [17], and flower orientation [18] influence the plant’s efficiency in pollen transfer or the pollinator’s efficiency in obtaining nectar.

Raup [19] proposed a general framework for discovering the functional consequences of morphological variation: the theoretical morphospace paradigm. This approach explores “n-dimensional geometric hyperspaces produced by systematically varying the parameter values of a geometric form” [20]. Raup devised a simple and elegant way to describe variation in the shape of mollusk shells using a 3-parameter equation, then tested the hydrodynamic performance of artificial shells fabricated to sample a wide range of the total 3-dimensional morphospace. Because the bounds of this theoretical morphospace are not constrained by naturally existing forms, it enables unbiased study over all the theoretically possible forms, which includes those that never have occurred in nature.

Given the phenomenal diversity of flower morphologies in nature, constructing a mathematical flower model capable of quantifying and easily manipulating morphological variation along multiple orthogonal axes (morphospace) is the logical first step in our effort to distinguish between mutualistic and antagonistic plant-pollinator interactions. Following the theoretical morphospace paradigm [20], we previously defined flower morphology with a 4-parameter equation (Fig. 1A), fabricated the flower models with a 3D printer (Fig. 1B), and experimentally tested a hawkmoth pollinator’s ability to exploit the flower’s nectar. We showed that both flower curvature and nectary diameter influence the hawkmoth pollinator’s performance [21].

**Figure 1.**
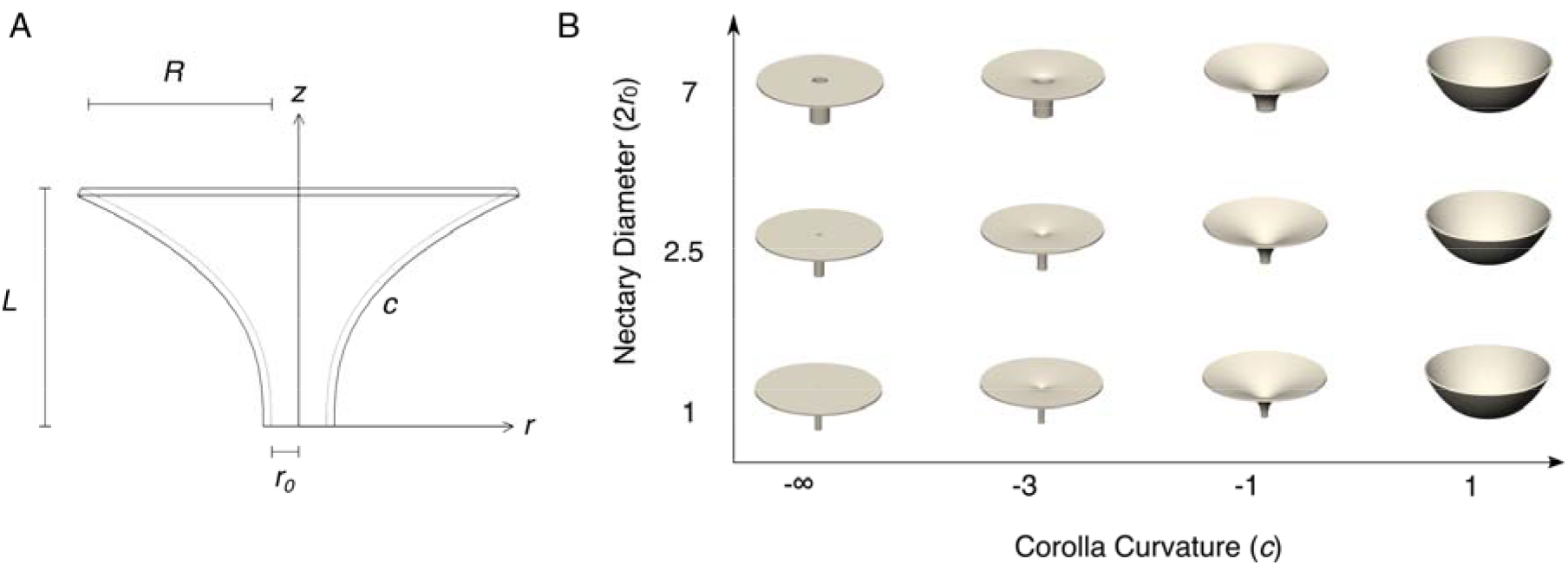
Design of artificial flowers. (A). Side view of a flower, showing the four parameters in the equation. (B). 3D rendering of some representative flowers with variation in nectary diameter and corolla curvature. The corolla tube length *L* is fixed at 20 mm and the overall diameter 2(*R* + *r_0_*) is fixed at 55 mm.

In this study, we took a two-stage approach to examine the potential conflict between plants and pollinators. First, we thoroughly explored the flower morphospace along two flower shape axes (corolla curvature and nectary diameter) to search for regions which might generate an evolutionary conflict of interest between plants and a hawkmoth pollinator. Then, we explicitly quantified the fitness of plants and hawkmoth pollinators in this critical region of morphospace by using a combination of machine vision techniques and electronic sensors to detect both the efficiency of nectar acquisition by the pollinator (a correlate of pollinator fitness) and the number of pollinator contacts with the plant’s reproductive parts (a proxy for plant fitness).

## Materials and Methods

### (A) Animals

Individual *Manduca sexta* hawkmoths were obtained from a colony maintained by the Department of Biology at the University of Washington. Hawkmoths were flower-naïve and unfed for 1-3 days post-eclosion. The hawkmoths used in foraging experiments were obtained haphazardly with respect to sex. Each hawkmoth’s sex, body weight, and proboscis length were recorded before it was used in experiments.

### (B) Fabrication of artificial flowers

The artificial flowers used in this study were fabricated in polyvinyl chloride (PVC) plastic using a 3D printer by the methods described in [21]. The printed flower model was white, rigid and effectively scentless. Flower models were designed in SolidWorks (Dassault Systèmes SolidWorks Corporation, Waltham, Massachusetts, USA) based on an equation with four flower shape parameters: corolla curvature, nectary radius, flower length, and corolla radius. In a manner similar to that of Raup [20], we used a parametric equation for a surface generating curve [21]:

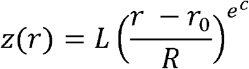

where *z* represents the longitudinal axis of the flower model, *r* represents the radial distance of the corolla from the central *z*-axis, *c* is a curvature parameter determining the lateral profile of the corolla, *r_0_* is the nectary radius, *L* is the flower length, and *R* is the lateral extent of the corolla from edge of the nectary to the outer lip of the flower (*i.e.*, the full radius from the central z-axis is equal to *r_0_* + *R*) (Fig. 1A). This curve is rotated about the z-axis and given a 1 mm thickness to create a 3D model.

### (C) First stage experiment to define the critical region of flower morphospace

#### a Experimental apparatus and flight arena

Artificial flowers were arranged in a 6 by 6 square array with flower centers spaced at 30.5 cm (Fig. S1). Modular T-slot extruded aluminum (80/20® Inc.) was used as the structural support for the flower array. Each 36-flower array was populated with 6 distinct flower morphologies, present at equal frequencies. Flower positions were randomized before each foraging trial. Each artificial flower’s nectar reservoir was filled with 20 μL of 20% w/v sucrose solution. The flower array was located inside a flight chamber (2.4 m width x 4.0 m length x 2.5 m height). Two dim white LED lights illuminated the arena from above, calibrated to a combined illuminance of 0.1 lux at flower level to simulate moonlight conditions. The flight chamber was also illuminated from above with a single infrared light (Magnalight LEDLB-16E-IR; 790-880 nm flat emission peak, modified by removing the light’s focusing lenses to create even, diffuse lighting), invisible to *M. sexta*, to allow video recording of the hawkmoth’s foraging trajectory (Fig. S1).

Each artificial flower was held in the array by a 3D-printed bracket that contained an infrared (IR) emitter-detector pair, creating an infrared beam sensor. When a hawkmoth’s proboscis entered the nectar reservoir of any artificial flower, the infrared beam was broken, and this event was recorded through a data acquisition system consisting of an Arduino microcontroller (Sparkfun: DEV-11021) and a laptop computer (Fig. S1). Data collection ended when all six flowers of any single morph were exploited. An infrared-sensitive video camera was mounted above the flower array facing straight down, to capture the hawkmoth’s flight trajectory as it foraged on the artificial flowers.

The air temperature of the flight arena was kept at approximately 24°C. To stimulate hawkmoth foraging, 5 μL of a 7-component synthetic scent mixture [21] was placed in the flight chamber 5 min before the experiment began. The synthetic scent mixture mimics the scents emitted by the flower of the hawkmoth-pollinated plant *Datura wrightii*.

#### b Experimental treatments and sample size

A foraging trial ended when the hawkmoth left the flower array and no longer visited flowers. Foraging trials typically lasted from 4 to 12 minutes. No hawkmoth participated in more than one foraging trial. After each trial, emptied flowers were counted, empty nectar tubes were replaced with fresh nectar tubes, and the positions of the flower morphs were re-randomized in preparation for the next foraging trial.

Hawkmoths were tested with various combinations of corolla curvature (*c*: –∞, −4, −3, −2, −1, 0, 0.375 and 1) and nectary diameter (2*r_0_*: 1, 1.75, 2.5, 3.25, 5, and 7 mm). All flowers had a length (*L*) of 20 mm and an overall diameter 2(*R* + *r_0_*) of 55 mm (Fig.1B). In addition to flower exploitation data in the form of counts of emptied flower morphs, the number of visits paid to each flower morph was also recorded by analyzing the video recording of each foraging trial. Exploitation and visitation data were collected for 125 foraging trials (*i.e.*, 125 different individual hawkmoths).

#### c Data analysis

Two-way analysis of variance (ANOVA) was used to test for differences in flower visitation frequency (an estimate of plant fitness, calculated as the number of times hawkmoths visit each flower morphology per trial, measured from the video captured for each foraging trial) and hawkmoth foraging success rate (an estimate of pollinator fitness, calculated as the number of emptied flowers per morphology per trial divided by the number of visits to that morphology per trial; flower morphologies which received 0 visits in a trial were excluded from foraging success rate analysis).

### (D) Second stage experiments focused on the critical region of flower morphospace

#### a Experimental apparatus and flight arena

In the second stage experiments, we focused on flower-hawkmoth interaction in the region of flower morphospace identified in the first stage as most sensitive to variation in corolla curvature (*c*) and nectary diameter (2*r_0_*). In the second stage experiments, individual hawkmoths were presented with only one nectar-bearing flower in the flight arena, to examine explicitly the effectiveness of each pollinator visitation event. The experimental flower apparatus consisted of a 3D-printed flower model, an “anther/stigma” sensor assemblage to record the physical contact between the pollinator and the “reproductive parts” of the flower, and a nectary assemblage to provide nectar and detect nectar level changes (Fig. 3B).

A small (4 x 4 x 1.45 mm) 3-axis accelerometer (ADXL335, SparkFun, Niwot, Colorado, USA) was used as a proxy for the flower’s reproductive organs (*i.e.*, anthers and stigma). The accelerometer was solder-connected with ultra-thin silver wire (0.14mm) and supported on a 20mm long thin stainless steel wire (0.13mm), mimicking the filament and style to support the anthers and stigma in real flowers. The other end of the steel wire was inserted into a rigid stainless steel tube to fix the free vibrating length of the wire, so that the natural frequency of the wire (about 18.4 Hz) did not change during the experiment. The accelerometer was positioned in the center of the corolla, 10 mm above the flower top plane.

A 25 mm long rigid plastic tube (inside diameter = 3.18mm) connected the printed flower and the nectary assemblage. The distance from the plane of the flower top to the bottom of nectar reservoir was 70mm, which is shorter than the average proboscis length of *M. sexta* (82.5 ± 3.25 mm, N = 58). The same infrared emitter-detector pair as in the first stage experiments was installed on the side of nectar base, allowing infrared light to pass through the bottom of the nectar reservoir to record nectar level changes. We constructed an automatic nectary refiller using a micro-injector pump, controlled by an Arduino Uno microcontroller, to push the nectar-filled Hamilton syringe with a stepper motor (adapted from [22]), to automatically replenish 20 μL of nectar during the trial. The micro-injector had enough volume capacity to refill the nectar reservoir 25 times.

The experimental flower apparatus was affixed to the floor of a flight chamber. Two distractor flowers (morphology: *r_0_* = 1 mm, *c* = −3, *L* = 30 mm, *R* = 24 mm), which had no accelerometer assemblage or nectar supply, were placed 25 cm on either side of the experimental flower. The distractor flowers were used to distract the hawkmoth from the experimental flower, so that the nectar in the experimental flower could be replenished after the hawkmoth left. A webcam (C170, Logitech, Lausanne, Switzerland) was affixed to the ceiling of the flight chamber, facing vertically down at the experimental flower. The videos were taken at a frequency of 5 frames/sec. The lighting and scent conditions in the flight chamber were identical with the first stage experiments (Fig. 3A).

#### b Experimental treatments and sample size

Moths were presented with one instrumented artificial flower per trial. We tested four different corolla curvatures (*c*: –∞, −3, −1, and 1; fixing other floral shape parameters at 2*r*_0_ = 3 mm; *L* = 30 mm; *R* = 23.5 mm). A 3 mm nectary diameter was used to account for the presence of the artificial anther/stigma (the diameter of the wiring is about 0.5 mm) partially occluding the corolla tube, leaving an open diameter of 2.5 mm. This diameter is consistent with measured nectary diameter values for hawkmoth-pollinated flowers such as *Petunia axillaris* and *Datura wrighttii* which have nectary diameters ranging from 1 to 2.5 mm [23]. Accelerometer and infrared nectar sensor data were collected through an Arduino Uno microcontroller at a frequency of 1 kHz. A foraging trial ended when either: 1) the nectar had been emptied 25 times (*i.e.*, reached the capacity of the micro-injector); or, 2) four minutes had passed since the last visit to the instrumented flower. For each flower morph, we collected data from 14-15 hawkmoths. The total number of hawkmoths over all trials was 58.

#### c Data analysis

The videos taken by the webcam were analyzed with a customized Python program (Github repository: https://github.com/foenpeng/Controller-of-experiment-platform.git). Because the instrumented flower and webcam were both static, the hawkmoth’s location in each frame can be obtained by frame subtraction. A reference frame was taken before a hawkmoth started foraging. Every new frame was subtracted from this reference frame. The centroid of the largest contour in every subtracted frame was used as an estimate of the hawkmoth’s position. The coordinates of the centroid were compared with a 125 mm radius circle (about the total length of a hawkmoth’s body plus a fully extended proboscis) centered on the experimental flower to determine the presence/absence of the hawkmoth pollinator. A pollinator visit was recorded whenever the hawkmoth centroid was inside the 125 mm radius.

In the second stage experiments we focused on measuring the pollinator’s visit *quality*, instead of visit *quantity* as in the first stage experiments. In the second stage experiments pollinator fitness was estimated as net rate of energy gain per visit, and plant fitness was estimated as hit counts to the artificial anther/stigma per visit. The plant’s and pollinator’s fitnesses (defined below) with respect to flower morphology (corolla curvature) variation were analyzed by ANOVA. The plant’s and pollinator’s fitnesses with respect to hawkmoth morphology (proboscis length) variation were analyzed by linear regression.

We used the amount of acquired energy (from nectar) divided by the total time spent within 125 mm of the instrumented flower during each visit to represent the pollinator’s net rate of energy gain per flower visit, which is a correlate of hawkmoth fitness [24]. The total amount of time that each hawkmoth spent during each visit to the instrumented flower was calculated from the video tracking data. The amount of energy acquired is fixed for every visit: the hawkmoth always consumes all of the 20 μL of 20% sucrose nectar in the container if it reaches nectar, which provides a maximum of 64.8 J of energy assuming all energy ingested is available; if it fails to reach nectar, it obtains 0 J energy. The rate of energy expenditure is minute relative to the rate of energy gain (2-3% of acquired energy [24]), so energy expenditure is neglected.

As a proxy for the plant’s fitness, the total number of “hits” on the accelerometer (representing the anthers and stigma of a real flower) during each hawkmoth visit was inferred from the accelerometer data. The analog signal readout from the accelerometer was calibrated using gravitational acceleration (*g* = 9.8 m s^-2^) when the accelerometer was in a static state. The total acceleration was calculated as the square root of the sum of squares of acceleration in each of the three axes (x, y, z). A “hit” was identified with a peak detection algorithm (Python peaktutils package), with total acceleration greater than 3 *g* counted as a hit.

## Results

### (A) The first stage experiment identifies a critical region in flower morphospace where the interests of plants and pollinators are both sensitive to corolla curvature variation

#### a Flowers with gentle curvatures are visited less frequently by hawkmoths

Flower curvature significantly influences the flower visitation frequency, the proxy for plant fitness in the first stage experiment (2-way ANOVA, p = 2.56x10^-6^). Flowers with more extreme curvatures (*c* = –∞ or *c* = 1) received more visits than the flowers with gentle curvatures (*c* = –1 or *c* = –2) (Fig. 2C). Nectary diameter (p = 0.31) and the interaction between nectary diameter and corolla curvature (p = 0.29) have no significant influence on flower visitation frequency.

**Figure 2.**
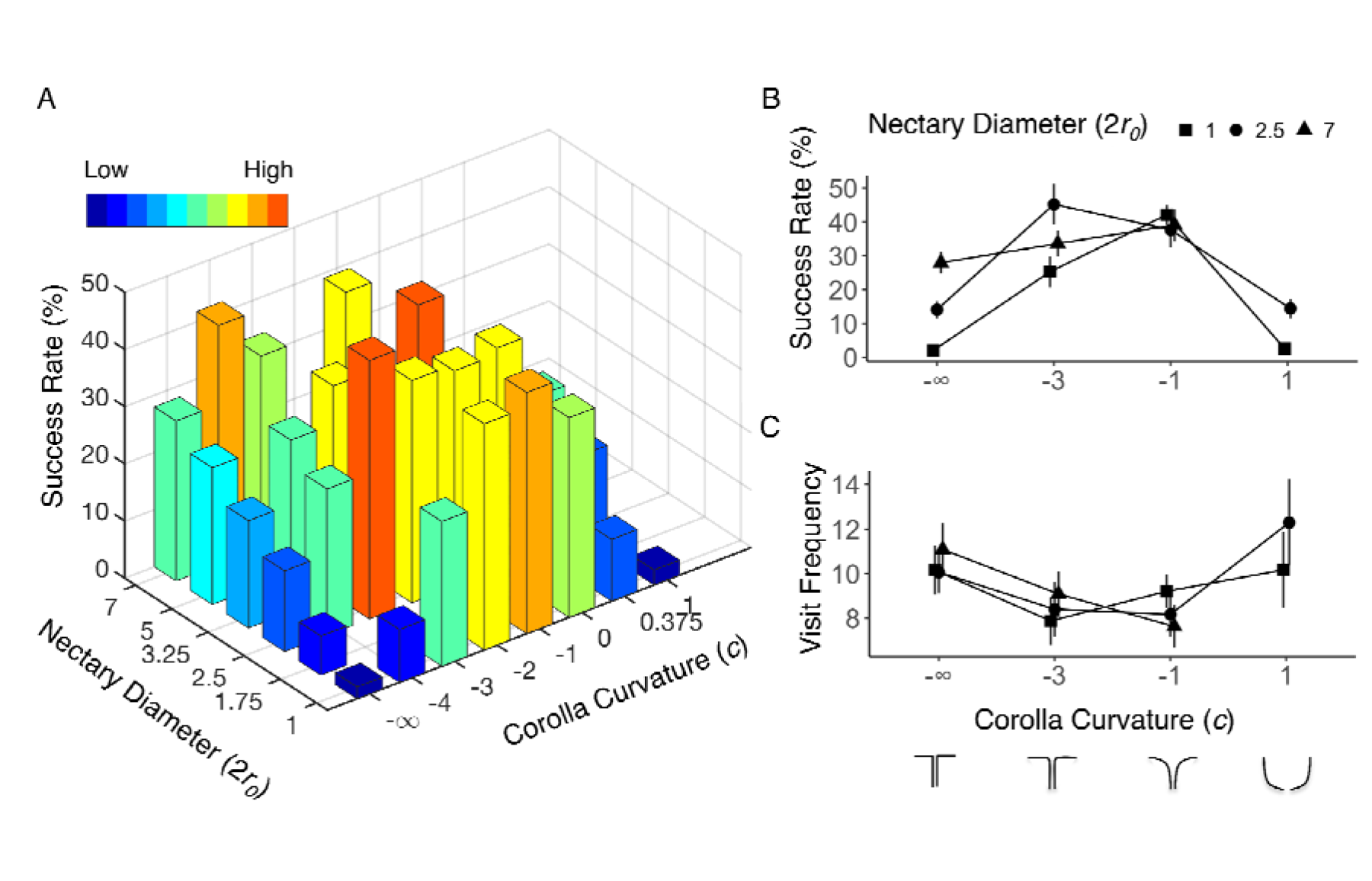
Hawkmoth foraging success rate and flower visitation frequency in the first stage experiment. (A). Hawkmoth foraging success rate (number of emptied flower/number of visits per trial, a correlate of pollinator fitness) across two dimensions of flower morphospace. Graphs B and C show some flowers that capture the range of variation along the two morphological axes from the full data set (Supplementary Table 1). (B). Hawkmoth foraging success rate. (C) Flower visitation frequency. Error bars represent± 1 standard error of the mean (SEM).

**Figure 3.**
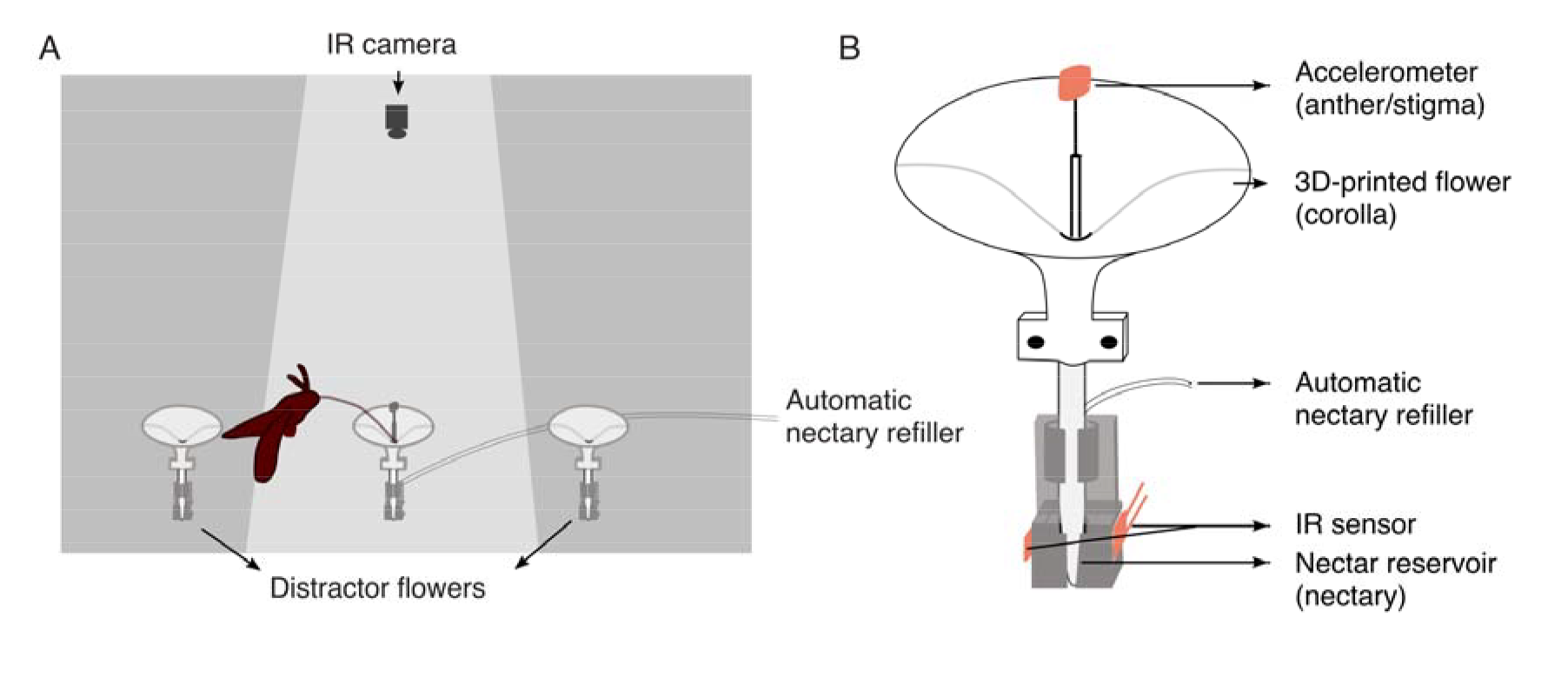
Second stage experimental setup. (A). Hawkmoth flight chamber. Only the center flower is instrumented and supplied with nectar. The flowers on the two sides are distractors. (B). Instrumented flower model, which is an enlarged view of the center flower in (A).

#### b Hawkmoth foraging success rate is maximized in gently-curved trumpet-shaped flowers

Foraging success rate, the proxy for pollinator fitness in the first stage experiment, is measured as the number of emptied flowers per morphology per trial divided by the total number of visits that the hawkmoth paid to that flower morphology. Two-way ANOVA shows that there is a significant effect on hawkmoth foraging success rate due to variation in nectary diameter (p = 7.18x10^-9^), corolla curvature (p = 2x10^-16^), and the interaction between nectary diameter and corolla curvature (p = 3.25x10^-5^).

The easiest flower morphology for hawkmoths to exploit (*c* = −3, 2*r_0_* = 2.5) yielded an average foraging success rate of 45% (± 6% SEM) while the most difficult one (*c* = -∞, 2*r_0_* =1) was fed upon with an average success rate of 2% (± 0.7% SEM) (Fig. 2A and 2B). A peak in foraging success occurs among trumpet-shaped flowers, with the hawkmoth’s performance decreasing steadily toward the extremes of corolla curvature: flat (*c* = -∞,) and bowl-shaped (*c* = 1) flowers (Fig. 2B). The difference in foraging success rate between trumpet-shaped flowers and flowers with extreme curvatures is magnified as the nectary diameter decreases (Fig. 2B).

#### c Hawkmoth foraging success rate is sensitive to curvature only when the nectary diameter is small

Individually, a large nectary diameter and a trumpet-shaped corolla curvature yield similarly high foraging success rates, at about 40% per foraging trial (Fig. 2B). Foraging performance decreases with the decrease of nectary diameter. However, poor foraging performance at a small nectary diameter (at or below 2.5 mm) can be rescued by trumpet-like curvature (*c* = −1 and −2) to maximal performance (Fig. 2A and 2B).

### (B) The second stage experiment reveals an evolutionary conflict of interest between the plant and the hawkmoth pollinator

#### a Variation in flower corolla curvature produces an evolutionary conflict

In the second stage experiment, nectary diameter was fixed at a dimension (2*r_0_* = 2.5 mm) that is relevant to naturally occurring hawkmoth-pollinated flowers [23] and also reveals the hawkmoth pollinator’s sensitivity to corolla curvature (Fig. 2B). To corroborate the conflict of interest revealed by a fitness estimate related to visit *quantity* in the first stage experiments, we designed our second stage experiment to characterize visit *quality*; *i.e.* effectiveness. The plant’s fitness was estimated as artificial anther/stigma hit counts per visit. The hawkmoth’s fitness was estimated as the net rate of energy gain per visit. In the second stage experiment, corolla curvature significantly influences both the plant’s (ANOVA, p = 1.92x10^-10^) and the pollinator’s fitness (ANOVA, p = 2x10^-16^).

The hawkmoth’s fitness is maximal for trumpet-shaped flowers (*c* = −1; 6.35 ± 0.22 J/s/visit SEM), and steadily decreases towards the extremes of flat (*c* = –∞; 1.73 ± 0.19 SEM) and bowl-shaped flowers (*c* = 1; 2.33 ± 0.17 SEM), producing a bell-shaped fitness curve (Fig. 4A). This result is concordant with the findings from the first stage experiment when we use foraging success rate as hawkmoth’s fitness measurement (Fig. 2B), and suggests that the influence of corolla curvature on hawkmoth’s foraging performance is robust to the differences in experimental design and fitness measurement between first stage and second stage experiments.

**Figure 4.**
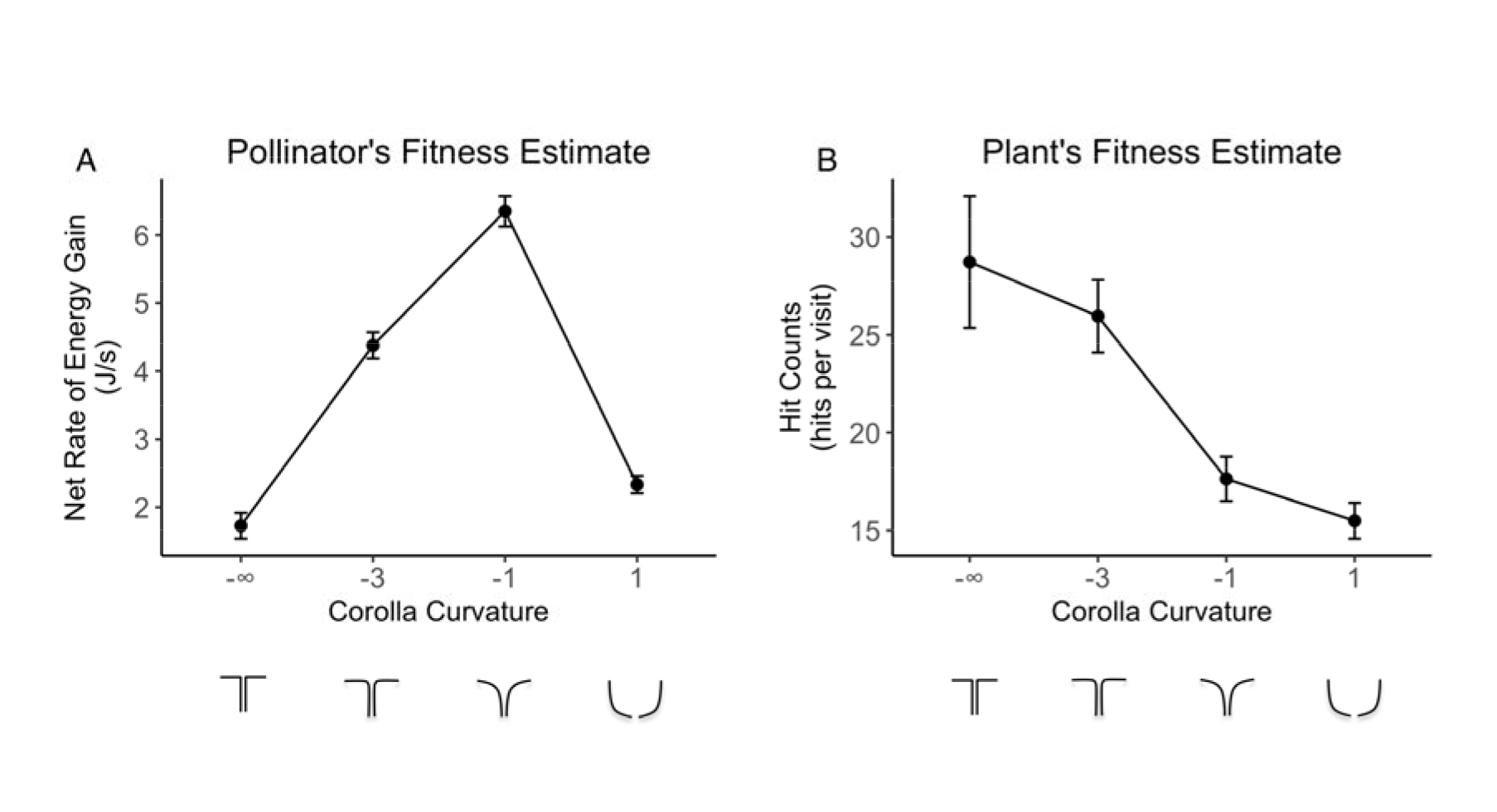
The fitness of pollinator and plant in the critical region of flower morphospace. (A). Hawkmoth pollinator’s fitness measured as net rate of energy gain; (B). Plant’s fitness measured as hit counts to the artificial anther/stigma. Error bars are ±1 SEM. Sample size (number of visits) of each flower morph is: c = −∞, N = 198; c = −3, N = 428; c = −1, N = 384; c = 1, N = 806.

The plant’s fitness estimate, based on the physical contacts between the hawkmoth’s body and the artificial anther/stigma (accelerometer), decreases steadily as corolla curvature increases (Fig. 4B). The bowl-shaped flower receives the fewest physical contacts (*c* = 1; 15.49 ± 0.91 hits/visit SEM), while the flat flower receives the most contacts (*c* = –∞; 28.73 ± 3.37 SEM) (Fig. 4B).

#### b Variation in hawkmoth proboscis length produces an evolutionary conflict

In addition to the engineering approach for testing for plant-pollinator conflict, we took advantage of the natural variation in hawkmoth proboscis length to evaluate its influence on plant and pollinator fitness. There is no significant difference (p = 0.51, two-tailed t-test) in proboscis length between male (82.19 ± 3.92 mm SD, N = 27) and female (82.77 ± 2.55 SD, N = 31) hawkmoths.

Hawkmoth proboscis length is positively correlated (p = 0.0001, F-test, Fig. S2A) with the hawkmoth’s fitness, but negatively correlated (p = 0.02, F-test, Fig. S2B) with the plant’s fitness.

## Discussion

Although traditionally the relationship between plants and their pollinators is viewed as mutualistic, they have different requirements from their interaction: efficient pollen transfer for the plant *vs*. efficient energy acquisition for the pollinator. Using a novel combination of engineering technologies (3D printing, artificial flowers instrumented with an IR nectar sensor and an accelerometer contact sensor, automated nectar replenishment, and machine vision) to explore this interaction in flower morphospace, we have shown that these disparate requirements generate a strong conflict of evolutionary interest (Fig. 2B-C, 4A-B).

As a proof-of-concept exploration, we first examined the hawkmoth pollinator’s performance in a two-dimensional flower morphospace, varying corolla curvature and nectary diameter. We showed that the influence of these two floral morphological features on pollinator performance is non-additive. Flowers with smaller nectary diameters are more difficult for the hawkmoths to exploit, but this effect could be countered by an appropriate corolla curvature (*e.g.* trumpet-shaped, *c* = −1) (Fig. 2A). The powerful morphospace paradigm [20] enabled us to study both the effects of single morphological traits and the interactions among them without bias or constraint, and we successfully demonstrated its utility by revealing a previously undiscovered interaction between corolla curvature and nectary diameter.

Corolla curvature is a key trait that significantly influences both flower visitation frequency and foraging success rates by the hawkmoth pollinator. We found that the flowers with more extreme curvatures, which are more difficult for the hawkmoths to exploit, were visited more frequently by hawkmoths (Fig. 2B-C). This result suggests that hawkmoths are unable to distinguish flower curvature differences at a distance (*e.g.*, visually), else they should preferentially visit those flower morphs that are easier to exploit, thus increasing their fitness. We suspect that when flowers are a limited resource for the pollinator, as they are in our experiments, hawkmoths make more attempts to visit the flowers whose nectar have not been emptied; *i.e*., they may remember the location of successfully exploited flowers and avoid re-visiting them. The flowers with more extreme curvatures are less likely to be emptied, and thus receive more visits. Since visit frequency is a component of the plant’s pollination success, this result suggests an evolutionary conflict of interest between plants and their hawkmoth pollinators with respect to flower visit *quantity*: although hawkmoths obtain less energy (*i.e.,* lower pollinator fitness) from the flowers with more difficult-to-exploit curvatures, those flowers receive more visits, thus potentially increasing pollination success (*i.e.,* higher plant fitness).

To further characterize this conflict of interest between plants and pollinators, our second stage experiment used a novel approach to measure visit *quality*; *i.e.* effectiveness in the critical region of flower morphospace where hawkmoth pollinator foraging performance is exquisitely sensitive to corolla curvature. Again, we found that there is a strong conflict of fitness interest between plants and pollinators with respect to corolla curvature when using a fitness measurement based on the quality of the relevant interaction: for the plant, contact with the flower’s reproductive organs (anther and stigma); and, for the pollinator, net rate of energy gain.

When corolla curvature is positive (bowl-shaped flower morphologies), the estimates of both the plant’s and the pollinator’s fitness are low. However, when corolla curvature is negative (trumpet to flat disc morphologies), there is a substantial conflict of interest. Although hawkmoth pollinators are less efficient at acquiring energy from the flowers with more extreme negative curvatures, the reproductive parts of those flowers receive more “hits”, which is a proxy for pollen transfer (Fig. 4A-B).

This result can be understood from the mechanical basis of the two disparate activities – nectar acquisition for the pollinator *vs*. pollen transfer for the plant. On the one hand, hawkmoths rely on mechanosensory information from the proboscis to locate the nectary in low light conditions [25]. Flower corolla curvature can act as a mechanical guide to assist hawkmoth pollinators in finding the nectary [21]. The flat disc flower (*c* = –∞) and bowl-shaped flower (*c* = 1) probably provide less passive guidance of the proboscis and more ambiguous mechanosensory information about the location of the nectary than does the trumpet shaped corolla (*c* = −1). On the other hand, to deal with the more abruptly changing curvature (*c* = −∞ and c = −3), hawkmoths must position their heads closer to the center of flower, which leads to stronger contacts with the plant’s reproductive parts (anther/stigma) (Supplementary Video S2 and S3). Gently curved flowers (c = −1) and bowl-shaped flowers (c = 1) have wider openings, with greater latitude for the position of the hawkmoth’s head, so the plant’s reproductive parts received fewer contacts. As a result, the intrinsic difference between the requirements of the plant and pollinator generate a strong conflict of evolutionary interest.

The conflict of evolutionary interests in corolla curvature motivated us to further explore a potential conflict by taking advantage of the natural morphological variation in the length of the hawkmoth pollinator’s proboscis. Hawkmoth proboscis length has long been suspected to play a key role in flower nectar spur evolution [13], but its effects on the fitness consequences on the plants and pollinators have seldom been explicitly tested (but see [26]). When presented with flowers having an invariant corolla tube length (70 mm, from the flower top plane to the nectar reservoir) shorter than the shortest hawkmoth *M. sexta* proboscis (72 mm) in our experiment, we found that hawkmoths with longer proboscides are better at nectar feeding, but make fewer contacts with the flower’s reproductive parts. The hawkmoth inserts its proboscis no further than necessary to obtain nectar [16]. A longer proboscis enables the hawkmoth to access the nectar resource with greater efficiency. However, the distance from the hawkmoth’s body to the center of the flower is also greater, so the hawkmoth’s body is less likely to contact the flower’s reproductive parts. As a result, a longer proboscis benefits the hawkmoth while harming the plant’s reproductive interests. Intuitively, a longer corolla tube or nectar spur on the flower would have the opposite effect on each party. This result corroborates previous field studies showing an evolutionary conflict between pollinator proboscis length and flower tube length [26,27]. It also lends support for the classical hypothesis that an arms race between plants and pollinators drives the evolution of long proboscides in hawkmoths and correspondingly long corolla tubes and nectar spurs in flowers [28]. Such an arms race can lead to the extreme morphologies typified by Darwin’s orchid and its hawkmoth pollinator, whose nectar spur and proboscis can be 30 cm or more in length [13].

It may be counterintuitive that the seemingly mutualistic plant-pollinator relationship could be masking underlying strong conflicts. However, there are several lines of evidence supporting the conflict hypothesis. First, conflicts of interest have been described in some obligate pollination systems, such as fig trees and fig wasps, and yucca and yucca moths [29]. Although those pollinators transfer pollen for their host plants, they also directly reproduce inside the hosts’ reproductive structure, and the larvae feed on the host plant’s seeds and inflorescences, which inevitably generates a fitness conflict. Second, the fossil record suggests that the earliest form of animal-vectored pollination might have evolved from pollen collecting behavior of pollen-eating insects – an antagonistic plant-herbivory interaction [1]. Nectar production by plants and nectar collection by pollinators may have evolved as a derived interaction to mitigate this strong conflict. Third, pollen deposition by pollinating animals is mostly involuntary. Cheaters exist widely in both plants and pollinators [30]. For example, 30% of orchid species have cheating strategies by either mimicking a rewarding flower (food deception) or mimicking female insects to attract naïve males (sexual deception) [31]. The legitimate pollinators of some plant species, such as hummingbirds and bumblebees, can also be nectar robbers, removing nectar through a hole pierced at the base of the flower [32] without providing pollination service to the plant.

The conflict hypothesis offers a new understanding of plant-pollinator coevolution. Notably, the rapid diversification of flowering plants – Darwin’s “abominable mystery” [33] – is better understood as the result of conflict rather than mutualism, because mutualistic interactions should be maintained by stabilizing selection, reducing diversification rates. In contrast, conflict of interest could promote rapid diversification in flower morphology. The divergent selective pressure between plants and pollinators resulting from their conflict of interest will increase variation in flower morphology within a plant species. Plant populations with highly variable flower morphologies could follow different evolutionary trajectories to occupy different plant fitness peaks by pollinator-mediated assortative mating among similar flower morphologies. Ultimately, this assortative mating could lead to plant speciation by pollinator shift – the origin of new plant species pollinated by different pollinator guilds (*e.g*., hawkmoths, bumblebees, hummingbirds, bats) best able to exploit each alternative flower morph. The ongoing conflict between plants and their pollinators produces a coevolutionary “arms race” that never reaches equilibrium, accounting for the observed rapid plant speciation over long periods of evolutionary time.

In accordance with our finding, most cases of rapid coevolutionary diversification involve conflict of interests. For example, predator-prey [34], host-parasite [35], and even males and females involved in sexual reproduction [12] generate evolutionary arms races. Novel strategies could be favored by the antagonistic parties to counter the adaptation to each other, promoting rapid evolution on both sides of the conflict. Our results suggest that the conflict of interest between plants and pollinators might also be a prevalent and general theme in the pollination interaction. Although we recognize the cooperative aspect of pollination relationship, in terms of nectar rewards and pollen transfer service, we argue that the hidden conflict of interests between the two parties, like the evolution of corolla curvature, are more likely to promote flowering plant diversification.

The combination of 3D printing technologies, electronic sensing, and machine vision has enabled us to rationally design flower morphologies, accurately generate flower models with desired parameters, and automate high-throughput behavioral data collection during plant-pollinator interactions. Field pollination experiments could also benefit from the deployment of such engineering technologies, especially for studying night-foraging pollinators, such as hawkmoths and bats [36].

We can extend our exploration of flower morphospace to other features (*e.g*., corolla tube length, petal number and shape, color, scent, texture), and also map the fitness landscapes of animals representing other pollinator guilds to see if the conflict found in this study is generalizable. If complemented with *in plastico* experimental evolution on artificial flower populations, we could further investigate – in real time – how the divergent selective force exerted by different pollinator guilds drives flower pollination syndrome divergence.

## Conclusion

In this study we employed a novel engineering approach to investigate a longstanding evolutionary mystery – the rapid radiation of flowering plants. Our results support the notion that a strong conflict of fitness interest exists between plant and pollinator, which could drive flower morphological diversification and contribute to plant speciation by pollinator shift.

## Data Accessibility

The datasets supporting this article have been uploaded as part of the supplementary material.

## Author’s Contributions

F. Peng designed and carried out the second stage experiment, analyzed the data, and drafted the manuscript; E. O. Campos designed and carried out the first stage experiment, and participated in data analysis and drafting the manuscript; J.G. Sullivan participated in the second stage experiment design and execution; N. Berry and B. B. Song participated in the first stage data collection; T. Daniel and H. D. Bradshaw, Jr. conceived of the study, participated in the experimental design, and edited the draft manuscript. All authors gave final approval for publication.

## Acknowledgments

We thank B. Nguyen for his expert care of hawkmoths, and T. Deora for help taking the high-speed video and drawing the experimental set-up diagram. Two undergraduate assistants, S. Wang and C. Fang, also contributed significantly to the data collection for the first stage experiment.

## References

1. Crane, P. R., Friis, E. M. & Pederson, K. R. 1995 The origin and early diversification of angiosperms. Nature 374, 27–33. (doi:10.1038/374027a0)

2. Grimaldi, D. 1999 The co-radiations of pollinating insects and angiosperms in the Cretaceous. Annals of the Missouri Botanical Garden 86, 373–406. (doi:10.2307/2666181)

3. van der Niet, T. & Johnson, S. D. 2012 Phylogenetic evidence for pollinator-driven diversification of angiosperms. Trends in Ecology & Evolution 27, 353–361. (doi:10.1016/j.tree.2012.02.002)

4. McLaughlin, R. N. & Malik, H. S. 2017 Genetic conflicts: the usual suspects and beyond. Journal of Experimental Biology 220, 6–17.

5. Hodges, S. A. 1995 The influence of nectar production on hawkmoth behavior, self-pollination, and seed production in *Mirabilis multiflora* (*Nyctaginaceae*). American Journal of Botany 82, 197–204. (doi:10.1002/j.1537-2197.1995.tb11488.x)

6. Vereecken, N. J., Dorchin, A., Dafni, A., Hoetling, S., Schulz, S. & Watts, S. 2013 A pollinators eye view of a shelter mimicry system. Ann. Bot. 111, 1155–1165. (doi:10.1093/aob/mct081)

7. Seymour, R. S., White, C. R. & Gibernan, M. 2003 Environmental biology: Heat reward for insect pollinators. Nature 426, 243–244. (doi:10.1038/426243a)

8. Hembry, D. H., Yoder, J. B. & Goodman, K. R. 2014 Coevolution and the diversification of life. Am Nat 184, 425–438. (doi:10.1086/677928)

9. Schluter, D. & McPhail, J. D. 1992 Ecological character displacement and speciation in sticklebacks. Am Nat 140, 85–108. (doi:10.1086/285404)

10. Ehrlich, P. R. & Raven, P. H. 1964 Butterflies and plants: a study in coevolution. Evolution 18, 586. (doi:10.2307/2406212)

11. Summers, K., McKeon, S., Sellars, J., Keusenkothen, M., Morris, J., Gloeckner, D., Pressley, C., Price, B. & Snow, H. 2003 Parasitic exploitation as an engine of diversity. Biol Rev Camb Philos Soc 78, 639–675. (doi:10.1017/S146479310300616X)

12. Bonduriansky, R. 2011 Sexual selection and conflict as engines of ecological diversification. Am Nat 178, 729–745. (doi:10.1086/662665)

13. Wasserthal, L. T. 1997 The pollinators of the Malagasy star orchids *Angraecum sesquipedale, A. sororium* and *A. compactum* and the evolution of extremely long spurs by pollinator shift. Plant Biology 110, 343–359. (doi:10.1111/j.1438-8677.1997.tb00650.x)

14. Conner, J. K., Sahli, H. F. & Karoly, K. 2009 Tests of adaptation: functional studies of pollen removal and estimates of natural selection on anther position in wild radish. Ann. Bot. 103, 1547–1556. (doi:10.1093/aob/mcp071)

15. Harder, L. D. & Barrett, S. C. H. 1993 Pollen removal from tristylous *Pontederia cordata*: effects of anther position and pollinator specialization. Ecology 74, 1059–1072. (doi:10.2307/1940476)

16. Nilsson, L. A. 1988 The evolution of flowers with deep corolla tubes. Nature 334, 147–149. (doi:10.1038/334147a0)

17. Muchhala, N. 2015 Adaptive trade-off in floral morphology mediates specialization for flowers pollinated by bats and hummingbirds. Am Nat 169, 494–504. (doi:10.1086/512047)

18. Ushimaru, A., Dohzono, I., Takami, Y. & Hyodo, F. 2009 Flower orientation enhances pollen transfer in bilaterally symmetrical flowers. Oecologia 160, 667–674. (doi:10.1007/s00442-009-1334-9)

19. Raup, D. M. 1966 Geometric analysis of shell coiling: general problems. Journal of Paleontology 40, 1178–1190. (doi:10.2307/1301992)

20. McGhee, G. R. 2006 The geometry of evolution. Cambridge: Cambridge University Press. (doi:10.1017/CBO9780511618369)

21. Campos, E. O., Bradshaw, H. D. & Daniel, T. L. 2015 Shape matters: corolla curvature improves nectar discovery in the hawkmoth *Manduca sexta*. Funct Ecol 29, 462–468. (doi:10.1111/1365-2435.12378)

22. Wijnen, B., Hunt, E. J., Anzalone, G. C. & Pearce, J. M. 2014 Open-source syringe pump library. PLoS ONE 9, e107216. (doi:10.1371/journal.pone.0107216)

23. Campos, E. O. 2017. *Plant-pollinator interactions in an ecological and evolutionary context: the promising role of 3D-printing technology and mathematical modeling* (Doctoral dissertation). Retrieved from WorldCat database. (OCLC No.: 1014344606)

24. Sprayberry, J. D. H. & Daniel, T. L. 2007 Flower tracking in hawkmoths: behavior and energetics. Journal of Experimental Biology 210, 37–45. (doi:10.1242/jeb.02616)

25. Goyret, J. & Raguso, R. A. 2006 The role of mechanosensory input in flower handling efficiency and learning by *Manduca sexta*. Journal of Experimental Biology 209, 1585–1593. (doi:10.1242/jeb.02169)

26. Pauw, A., Stofberg, J. & Waterman, R. J. 2009 Flies and flowers in Darwin’s race. Evolution 63, 268–279. (doi:10.1111/j.1558-5646.2008.00547.x)

27. Anderson, B. & Johnson, S. D. 2008 The geographical mosaic of coevolution in a plant-pollinator mutualism. Evolution 62, 220–225. (doi:10.1111/j.1558-5646.2007.00275.x)

28. Micheneau, C., Johnson, S. D. & Fay, M. F. 2009 Orchid pollination: from Darwin to the present day. Botanical Journal of the Linnean Society 161, 1–19. (doi:10.1111/j.1095-8339.2009.00995.x)

29. Dufay, M. & Anstett, M.C. 2003 Conflicts between plants and pollinators that reproduce within inflorescences: evolutionary variations on a theme. Oikos 100, 3–14. (doi:10.1034/j.1600-0706.2003.12053.x)

30. Ghoul, M., Griffin, A. S. & West, S. A. 2014 Toward an evolutionary definition of cheating. Evolution 68, 318–331. (doi:10.1111/evo.12266)

31. Selosse, M.A. 2014 The latest news from biological interactions in orchids: in love, head to toe. New Phytologist 202, 337–340. (doi:10.1111/nph.12769)

32. Irwin, R. E., Bronstein, J. L., Manson, J. S. & Richardson, L. 2010 Nectar robbing: ecological and evolutionary perspectives. Annu. Rev. Ecol. Evol. Syst. 41, 271–292. (doi:10.1146/annurev.ecolsys.110308.120330)

33. Friedman, W. E. 2009 The meaning of Darwin’s ‘abominable mystery’. American Journal of Botany 96, 5–21. (doi:10.3732/ajb.0800150)

34. Geffeney, S., Ruben, P. C. & Brodie, E. D. 2002 Mechanisms of adaptation in a predator-prey arms race: TTX-resistant sodium channels. Science 297, 1336–1339. (doi:10.1126/science.1074310)

35. Spottiswoode, C. N. & Stevens, M. 2012 Host-parasite arms races and rapid changes in bird egg appearance. Am Nat 179, 633–648. (doi:10.1086/665031)

36. Nachev, V., Stich, K. P., Winter, C., Bond, A., Kamil, A. & Winter, Y. 2017 Cognition-mediated evolution of low-quality floral nectars. Science 355, 75–78. (doi:10.1126/science.aah4219)

